# The N’ terminus of Alpha-1 Giardin, a parasitic annexin orthologue, is essential for oligomerization and lipid-binding activity

**DOI:** 10.64898/2026.01.09.697555

**Authors:** Dawid Warmus, Corina D. Wirdnam, Erina A. Balmer, Tendai Muronzi, Manfred Heller, Özlem Tastan Bishop, Carmen Faso

## Abstract

Annexins are Ca²⁺- and membrane-binding proteins conserved across eukaryotes and involved in diverse membrane-associated processes, including unconventional protein secretion (UPS). In the intestinal parasite Giardia lamblia, annexins are represented by a family of 21 proteins termed alpha-giardins (αGs), all of which lack classical signal peptides yet are detected at the parasite surface and in the secretome. Here, we present a comprehensive functional analysis of αGs, with a particular focus on α1-giardin (α1G), a leading vaccine candidate against giardiasis and a putative virulence-associated UPS substrate. We show that αGs localize to peripheral endocytic compartments (PECs), specialized endo-lysosomal organelles implicated in protein release, and engage in multiprotein complexes enriched in UPS substrates. Using lipid-binding assays, cross-linking mass spectrometry, and structural modelling, we define conserved and divergent features of αG membrane interactions. Focusing on α1G, we combine site-directed mutagenesis, confocal microscopy, and mass photometry to demonstrate a critical role for its short N-terminal region in oligomerization and membrane association. Integration of these data with molecular dynamics simulations reveals how α1G oligomeric state and membrane occupancy are regulated at PEC membranes. Together, our findings establish a mechanistic framework for annexin-mediated UPS in Giardia and provide structural insights relevant to parasite virulence and vaccine development.

## Introduction

Annexins are a protein family found in almost every eukaryote investigated to date, including animals, plants, and fungi (Gerke and Moss, 2002a; Gerke *et al*., 2024; Warmus, Balmer and Faso, 2025). The basic functions of annexins revolve around their interaction with membranes and calcium ions (Liemann and ANITA LEWIT-BENTLEY, 1995; König and Gerke, 2000; Moss and Morgan, 2004; Rescher and Gerke, 2004) in the context of several cellular processes, including but not limited to membrane tethering (Mo *et al*., 2003), trafficking and organization of vesicles (Tcatchoff *et al*., 2012; Wang *et al*., 2016), ion transport (Huber *et al*., 1990; Berendes *et al*., 1993; Isas *et al*., 2000; Golczak *et al*., 2001), plasma membrane repair (McNeil *et al*., 2006; Bouter *et al*., 2011a; Boye and Nylandsted, 2016; Sønder *et al*., 2019) and unconventional protein secretion (UPS) (Popa, Stewart and Moreau, 2018; Balmer, Wirdnam and Faso, 2023).

Annexins are characterized by a 70 amino acid sequence known as the annexin repeat (or fold) (Gerke and Moss, 2002a; Moss and Morgan, 2004; Rescher *et al*., 2020), each repeat presenting five alpha-helixes, four of which are parallel to each other and one perpendicular. Four to ten annexin repeats constitute the structurally conserved core region. Together, these form a curved shape with concave and convex sides (Figure 1). The convex side is responsible for interaction with Ca^2+^ and lipids (Bitto *et al*., 2000; Gerke and Moss, 2002a) while the concave side, involving the N-terminal domain, takes part in protein complex formation (Gerke and Moss, 2002a; Rescher and Gerke, 2004). In contrast to the core region, the N-terminal head region exhibits significant variability and a generally disordered structure which determine a given annexin’s function and protein interaction partners.

**Figure 1.**
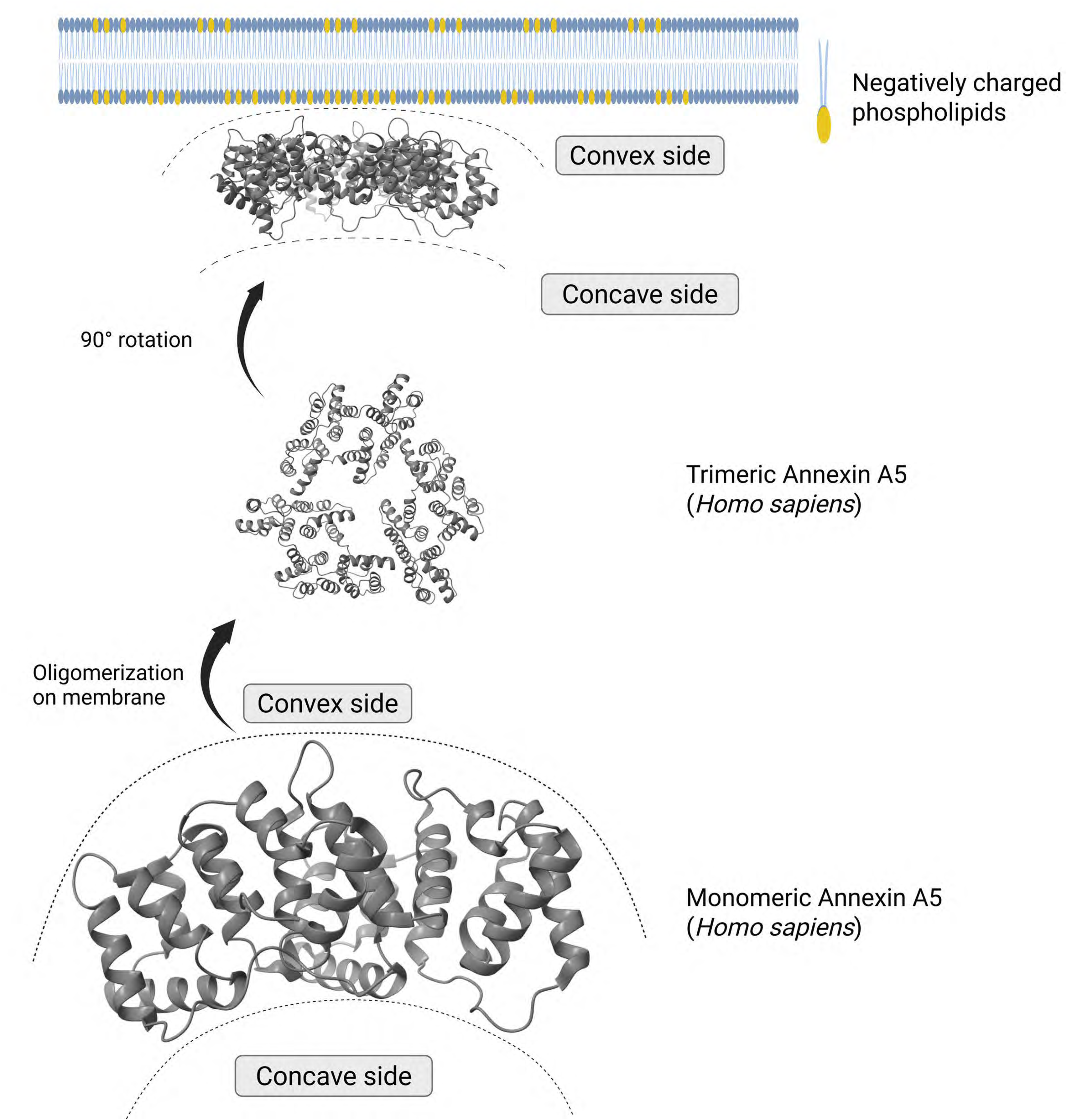
Human annexin A5 (anxA5) as a model for oligomerization and lipid binding at membranes. anxA5 is a well-characterized annexin, known to oligomerize prior to binding negatively charged lipids. As a general model for annexins, the convex side is responsible for contacting the membrane and is enriched for lipid binding residues.

At membranes, annexins can form various structures (Figure 1), ranging from simple aggregates to complex networks, depending on the specific annexin and membrane conditions (Oling, Bergsma-Schutter and Brisson, 2001; Mo *et al*., 2003; Ayala-Sanmartin *et al*., 2008; Grill *et al*., 2018). The specific mechanism of membrane binding varies among different annexins and is influenced by factors such as membrane lipid composition, presence of Ca^2+^ ions and pH (Langen *et al*., 1998; Gerke and Moss, 2002a; Moss and Morgan, 2004; Ladokhin and Haigler, 2005). Following membrane binding, annexins have been reported to insert themselves into the target membrane (Golczak *et al*., 2001; Gerke and Moss, 2002a; Mortimer *et al*., 2008). This process is thought to involve a conformational change in annexin structure, leading to lipid bilayer destabilization.

Annexins from the intestinal protist parasite *Giardia lamblia* (syn. *intestinalis, duodenalis*) are called alpha-giardins (αGs) (Bauer *et al*., 1999; Gerke and Moss, 2002b; Szkodowska *et al*., 2002; Elmendorf, Dawson and McCaffery, 2003; Weiland *et al*., 2003a; Vahrmann *et al*., 2008; Wei *et al*., 2010a; Weeratunga *et al*., 2012a; Einarsson *et al*., 2016a; Warmus, Balmer and Faso, 2025).Two members of the 21-strong protein family, alpha 1-giardin and alpha 11-giardin (α1G and α11G) were previously identified as putative UPS substrates following their detection in the surface proteome (Davids *et al*., 2019) and secretome (Ma’ayeh *et al*., 2017; Dubourg *et al*., 2018) of axenic Giardia cultures. Both proteins carry no *bona fide* signal sequence for secretion (Ma’ayeh *et al*., 2017; Dubourg *et al*., 2018; Davids *et al*., 2019). A recent investigation localized α1G and α11G to the parasite’s peripheral endocytic compartments (Balmer, Wirdnam and Faso, 2023) (PECs, formerly known as peripheral vacuoles), single membrane-bound endo-lysosomal organelles (Cernikova, Faso and Hehl, 2018, 2020; Faso and Hehl, 2019; Santos *et al*., 2022), previously implicated in protein release (Midlej, de Souza and Benchimol, 2019; Moyano *et al*., 2019; Balmer, Wirdnam and Faso, 2023). Furthermore, PECs-associated α1G and α11G participate in a protein complex which includes other UPS substrates, some of them recognized virulence factors in Giardia (Balmer, Wirdnam and Faso, 2023). Importantly, α1G is currently considered one of the strongest candidates for vaccine development for Giardiasis (Weiland *et al*., 2003b; Feliziani *et al*., 2011; Lopez-Romero *et al*., 2015; Davids *et al*., 2019), a worldwide water-borne diarrheal disease widespread in areas where water sanitation is problematic and safe drinking water cannot be guaranteed.

In this report, we present a broad functional characterization of αGs in terms of their subcellular location in the Giardia system, their lipid-binding behaviour, and their protein interactomes and predicted configurations using XL-MS (Liu *et al*., 2015; Klykov *et al*., 2018). Given its importance as a vaccine candidate for Giardiasis and its implication in virulence-related UPS mechanisms in Giardia, we narrow our focus to α1G. Using site-directed mutagenesis coupled to confocal microscopy and mass photometry (MP), we dissect the contribution of the short (less than five residues) N-terminal region of α1G and investigate oligomeric forms of both wildtype and mutant α1G variants in solution. These data are then used to inform simulations of molecular dynamics of α1G variants, to predict the behaviour of α1G variants at PEC membranes in terms of oligomeric stability and occupancy, and to develop a new working model for α1G function at PEC membranes.

## Results

### A subset of αGs localizes in close proximity to peripheral endocytic compartments

To determine where every αG accumulates within the Giardia cell, and especially to discover additional PECs-associated ones which might also be implicated in virulence-associated UPS, we generated constructs for the epitope-tagged expression of every member of the αG family. Despite attempts at expressing all αGs transgenically, only 15 of the 21 ORFs could be visualized as HA-tagged variants. Aside from α1G and α11G, previously shown to accumulate at PECs (Balmer, Wirdnam and Faso, 2023), an additional five ORFs (α5G, α6G, α12G, α15G and α17G) were also found in close proximity to these organelles (Figure 2). Several αGs presented a diffused and generally cytosolic pattern of accumulation, while others appeared distinctly at flagella (Figure 2).

**Figure 2.**
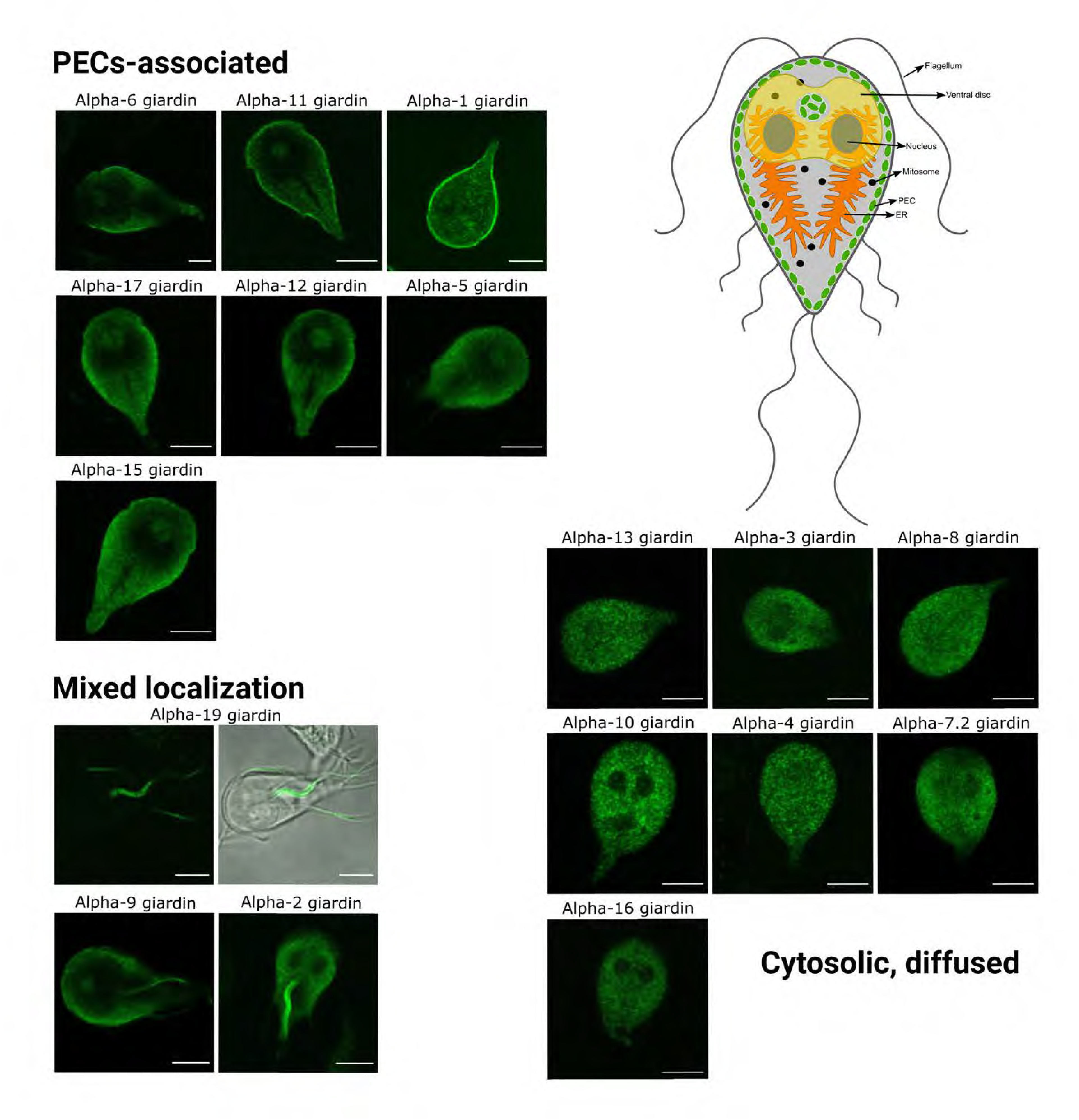
Confocal fluorescent light microscopy investigation of subcellular localization of epitope-tagged alpha-Giardins. PECs-associated: α1G, α5G, α6G, α11G, α12G, α15G and α17G. Cytosolic-diffused: α3G, α4G, α7.2G, α8G, α10G, α13G, α16G. Mixed: α2G, α9G and α19G. Schematic of a *Giardia* lamblia trophozoite, highlighting known compartments and PECs, shown in green.

### Lipid binding activity of Giardia-extracted PECs-associated αG complexes and native co-IP analysis

To determine whether differentially localized αGs present distinct lipid-binding behaviour, and based on confocal microscopy data, we selected four lines expressing PECs-associated αGs (6, 11, 2 and 1), and five expressing cytosolic-diffused αGs (8, 7.2, 9, 13, 16 and 19). Lines were chosen based on comparable levels of epitope-tagged variant expression, thereby excluding the impact of overall protein production on testing for lipid-binding capacity/specificity.

Native co-IP using HA-tagged αGs as affinity handles was followed by incubation of membrane lipid strips with the co-IP-ed eluate fraction. Antibody-based detection of the HA tag showed that, in contrast to αG complexes extracted from lines expressing epitope-tagged αGs 7.2, 9, 13, 16 and 19, where no appreciable signal was recorded, lipid residue binding was detected for protein complexes containing PECs-associated αGs (Figure 3). All tested IP-ed protein complexes containing PECs-associated αGs show affinity for selected PIP residues.

**Figure 3.**
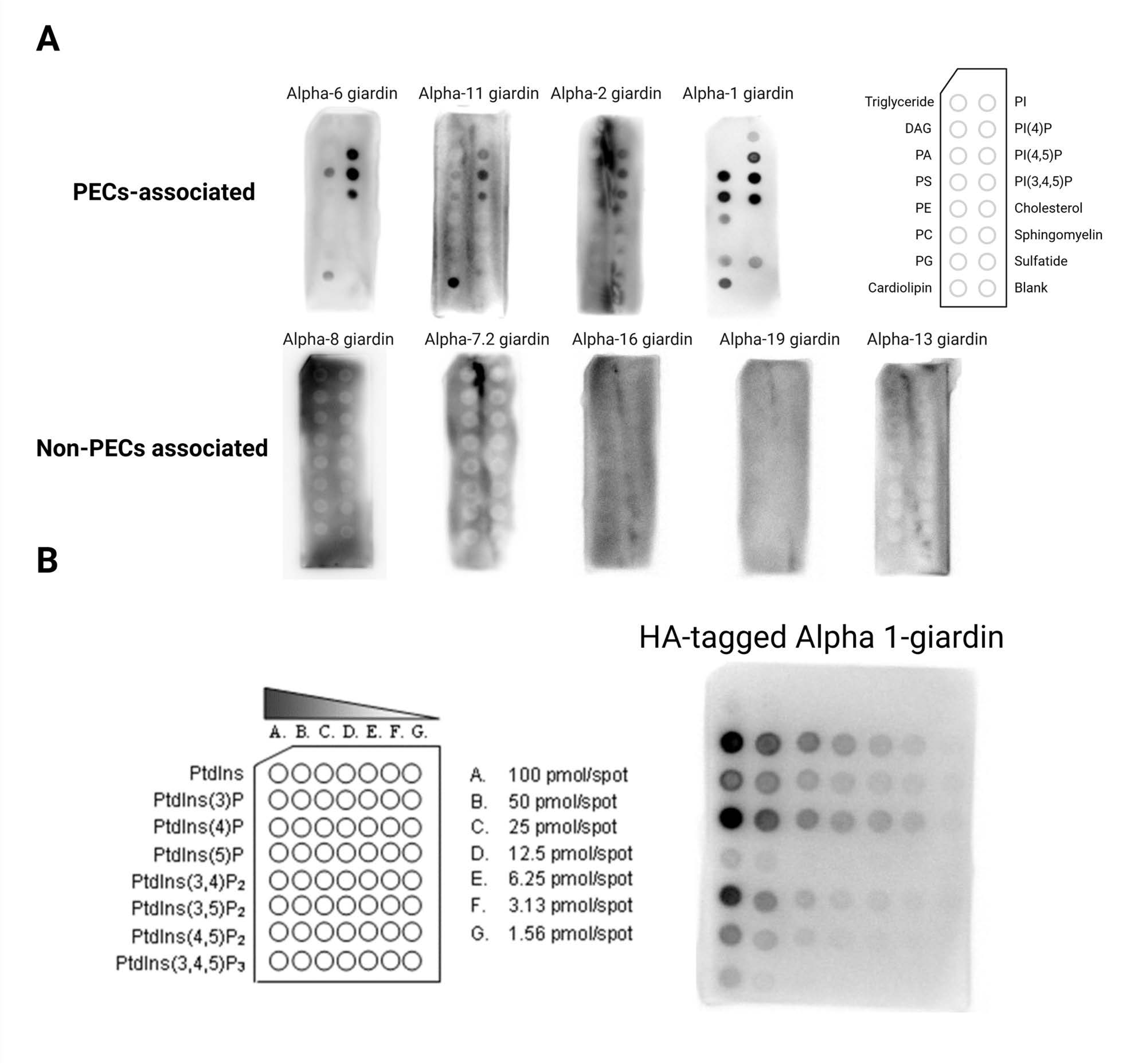
Lipid-binding activity of PECs-associated complexes IP-ed using epitope-tagged αGs as affinity handles. **A.** Protein complexes extracted from lines expressing PECs-associated αGs show a distinct binding profile using lipid strips, with PIP residues at the core, whereas lipid binding activity for complexes derived from lines expressing cytosolic/diffused αGs is not detected in the same experimental conditions. **B.** A finer analysis of lipid binding activity for α1G, with a focus on selected PIP variants using a PIP array.

To further investigate the composition of PECs-associated αG complexes, especially in comparison to α8G complexes which are never detected at PECs and show no appreciable lipid binding activity, biological replicates of co-IP-ed eluate fractions from lines expressing HA-tagged α1G (G25), α6G (G183), α11G (G190), α8G (G196), were analysed by tandem MS and compared to control fractions prepared from non-transgenic WB cells. A qualitative and comparative analysis using %riBAQ values across all samples (Figure 4A; Supplementary Data 1), followed by a ranking of the top 10 identified proteins in each sample (Figure 4B; Supplementary Data 1), shows a clear enrichment for each affinity handle compared to the WB control. Core interactomes for α1G, α6G and α11G-containing complexes are enriched for proteins previously shown to reside at PECs, including NEK kinases and UPS substrates with suspected roles in virulence (Xaa-dipeptidase/prolidase) (Balmer, Wirdnam and Faso, 2023). In contrast, the core interactome of α8G-containing complexes is consistent with its mainly cytosolic deposition and non-detectable lipid binding activity.

**Figure 4.**
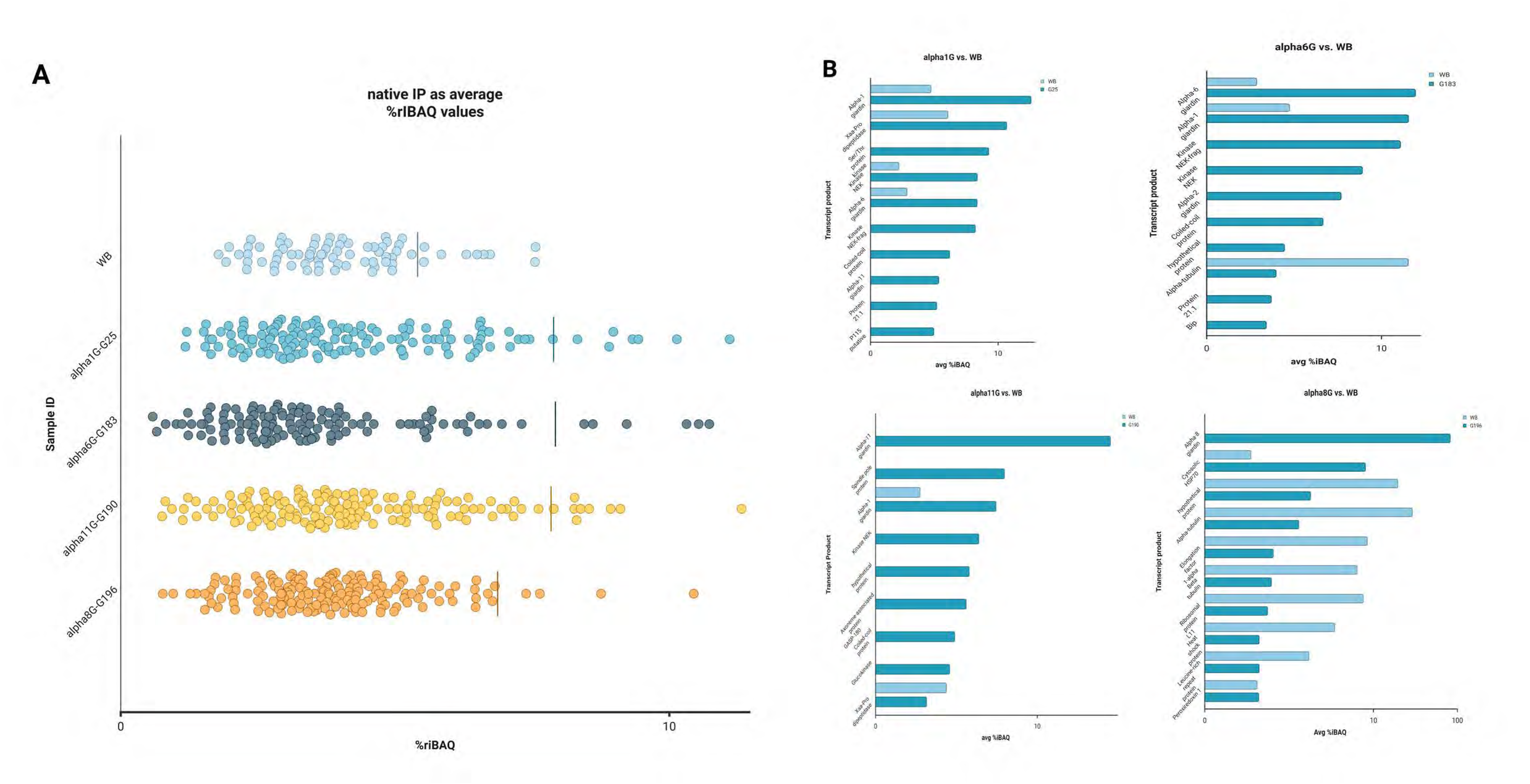
Native co-IP data analysis expressed as %rIBAQ values for HA-tagged affinity handles α1G, α6G, α11G, and α8G. **A.** Base-10 distribution of native co-IP data for each affinity handle, compared to native co-IP data for a non-transgenic WB control sample. **B.** Top protein hits based on %rIBAQ values for each native co-IP, compared to data for native co-IP from a non-transgenic WB sample.

XL-MS detection of inter- and intra-links in α1G complexes informs on structural configurations and possible sites of protein-protein interaction

To measure and calculate possible configurations of α1G-containing complexes, co-IP-ed eluate fractions from cells expressing HA-tagged α1G (G25), α6G (G183), α11G (G190), α8G (G196), were further analyzed by XL-MS using DSSO treatment followed by detection of covalently-bound peptides (Figure 5), either from within the same protein (<u>intra</u>-links) or between distinct proteins (<u>inter</u>-links), within the isolated complex (Liu *et al*., 2015) (Figure 5 and Supplementary Data 2).

**Figure 5.**
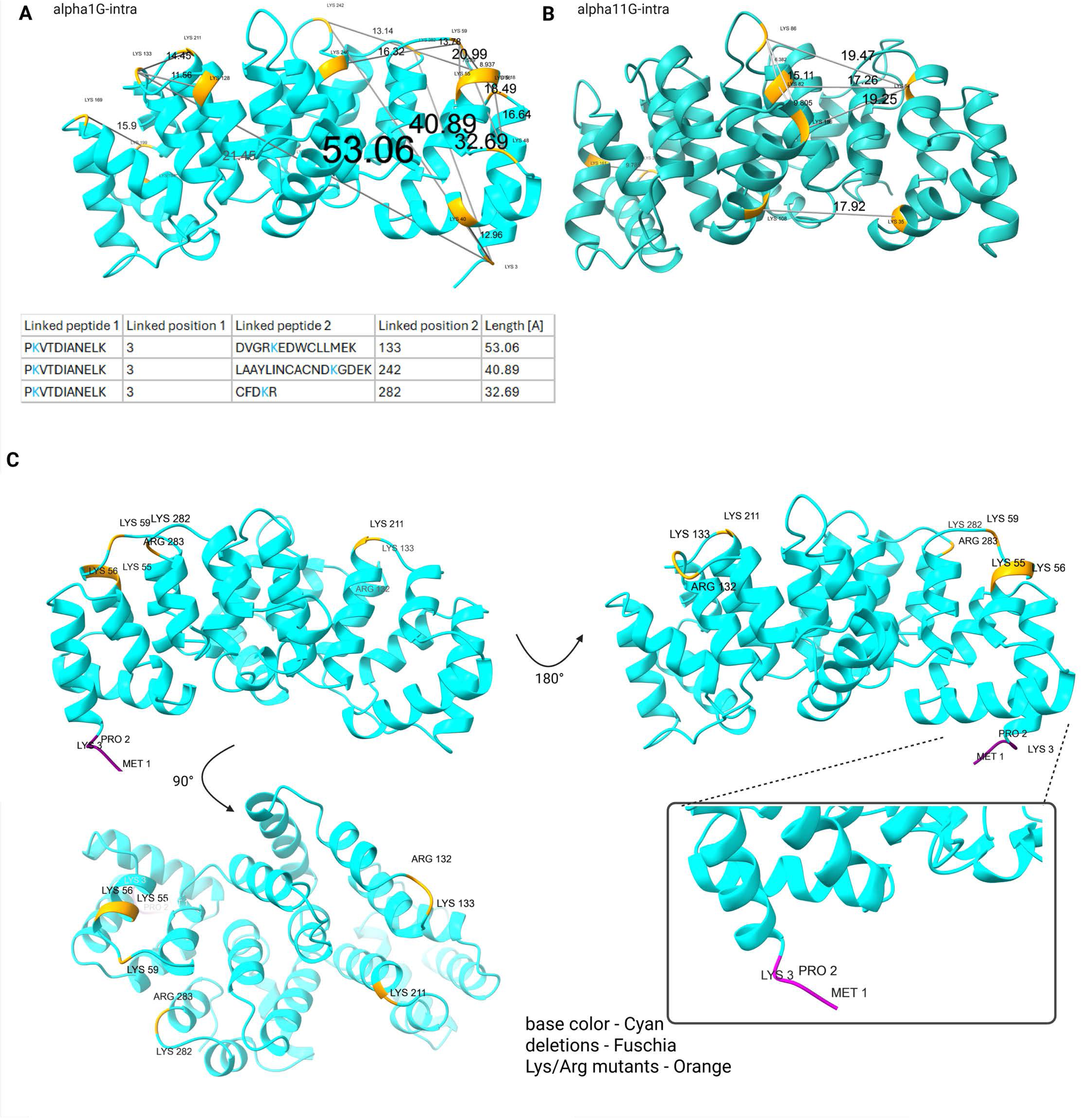
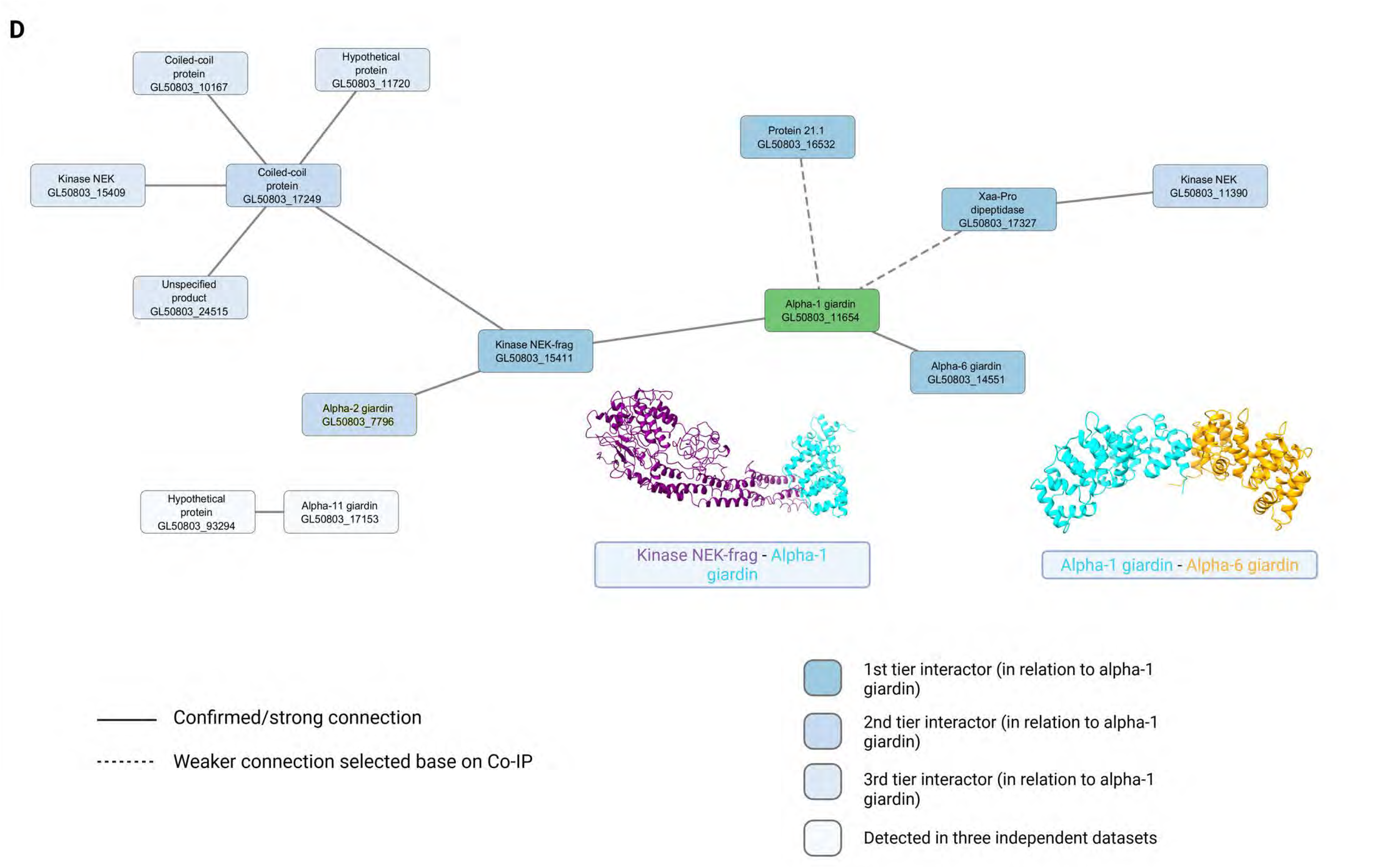
Structural modelling of inter and intra -links, following analysis of XL-MS data for α1G and α11G. **A.** XL-MS data analysis highlights the presence of intra-links compatible with oligomerization of α1G, mediated by the N terminal region and reported in detail in the table. **B.** XL-MS data analysis for α11G does not detect the presence of intra-links exceeding molecular interface crosslinking threshold. **C.** Mapping of selected residues and structural modelling analysis of α1G highlights the short N terminal region (fuschia) and putative lipid binding residues (orange). **D.** Visualization of tiered native co-IP and XL-MS data with a focus on α1G.

The detection of *intra*-links derived from covalently bound peptides from within α1G presents a discrepancy in terms of molecular interaction distances. The presence of intra-links significantly longer than the 25Å threshold (Figure 5A) can only be explained as derived from α1G homoligomers. In contrast, intra-links derived from within α11G (Figure 5B), α6G and α8G, never exceed the upper limit of 25Å, indicating how these αGs are unlikely to form oligomers (Supplementary Data 2).

The *inter*-links data (Figure 5D; Supplementary Data 2) point to a direct physical connection between α1G, a NEK kinase (ORF GL50803_15411), α6G and Xaa-dipeptidase. Surprisingly, although robustly present in all co-IP datasets previously reported (Balmer, Wirdnam and Faso, 2023) and presented here, neither α2G nor α11G are detected as being within molecular interaction distance (*ca*. 25Å) to α1G.

To further investigate, we mapped the exact residues deduced from both inter- and intra-links data for α1G, to identify interaction sites within the predicted structure of α1G which could be important for its function and structural configuration. This analysis led to the realization that the N-terminus of this protein is a hub for interactions, and that several positively charged residues on the convex side of the predicted structure could be important for lipid binding activity (Figure 5C).

### XL-MS informed targeted mutagenesis and subcellular localization of alpha1-giardin mutants and variants

To investigate the N-terminus of α1G and specific net positive charge residues for possible a role in protein localization, functionality and lipid-binding activity, as well as testing the predictive power of *in silico* structural analysis of the XL-MS data, we generated several constructs for expression of α1G mutant and deletion variants in transgenic Giardia lines (Figure 6).

**Figure 6.**
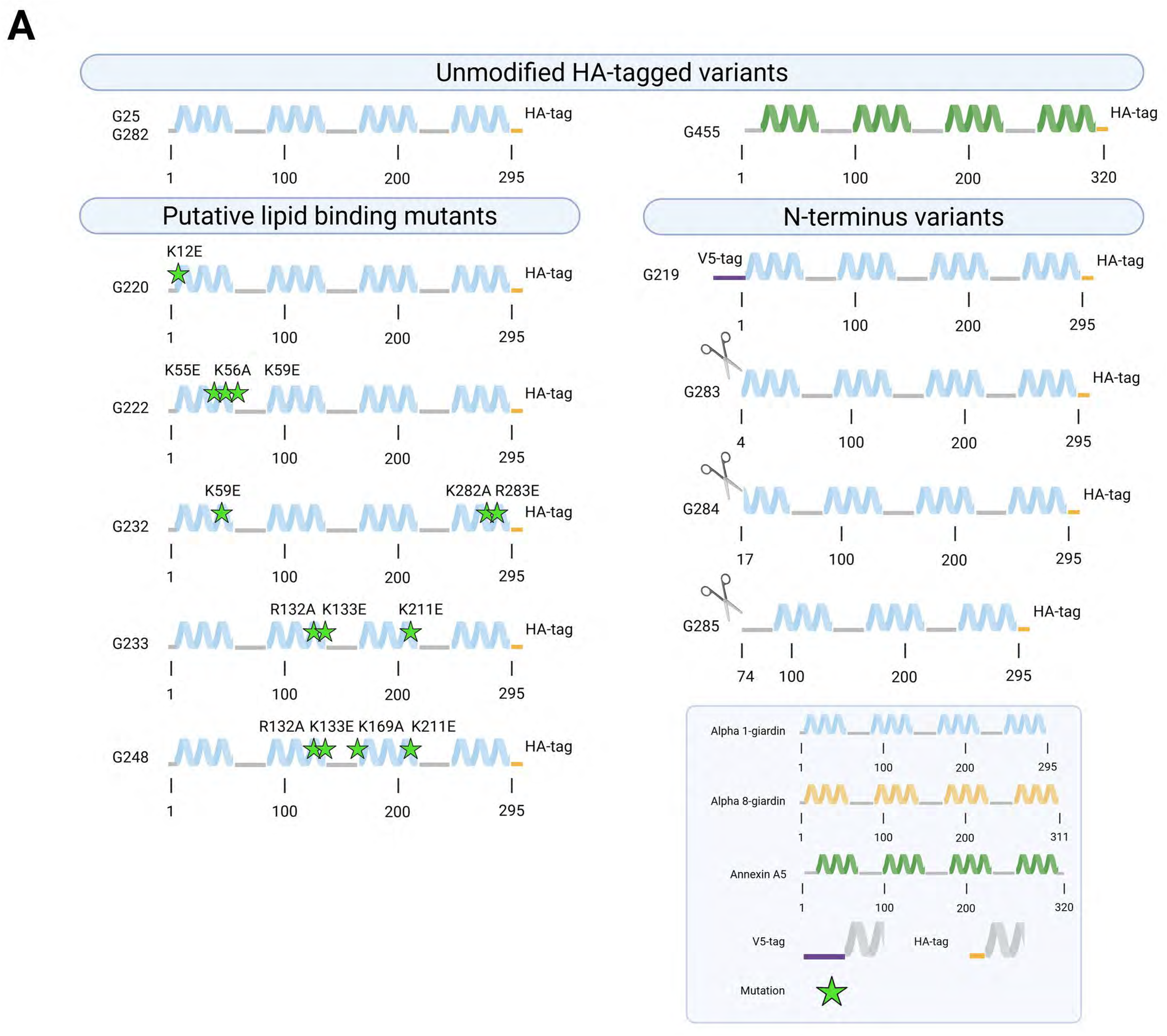
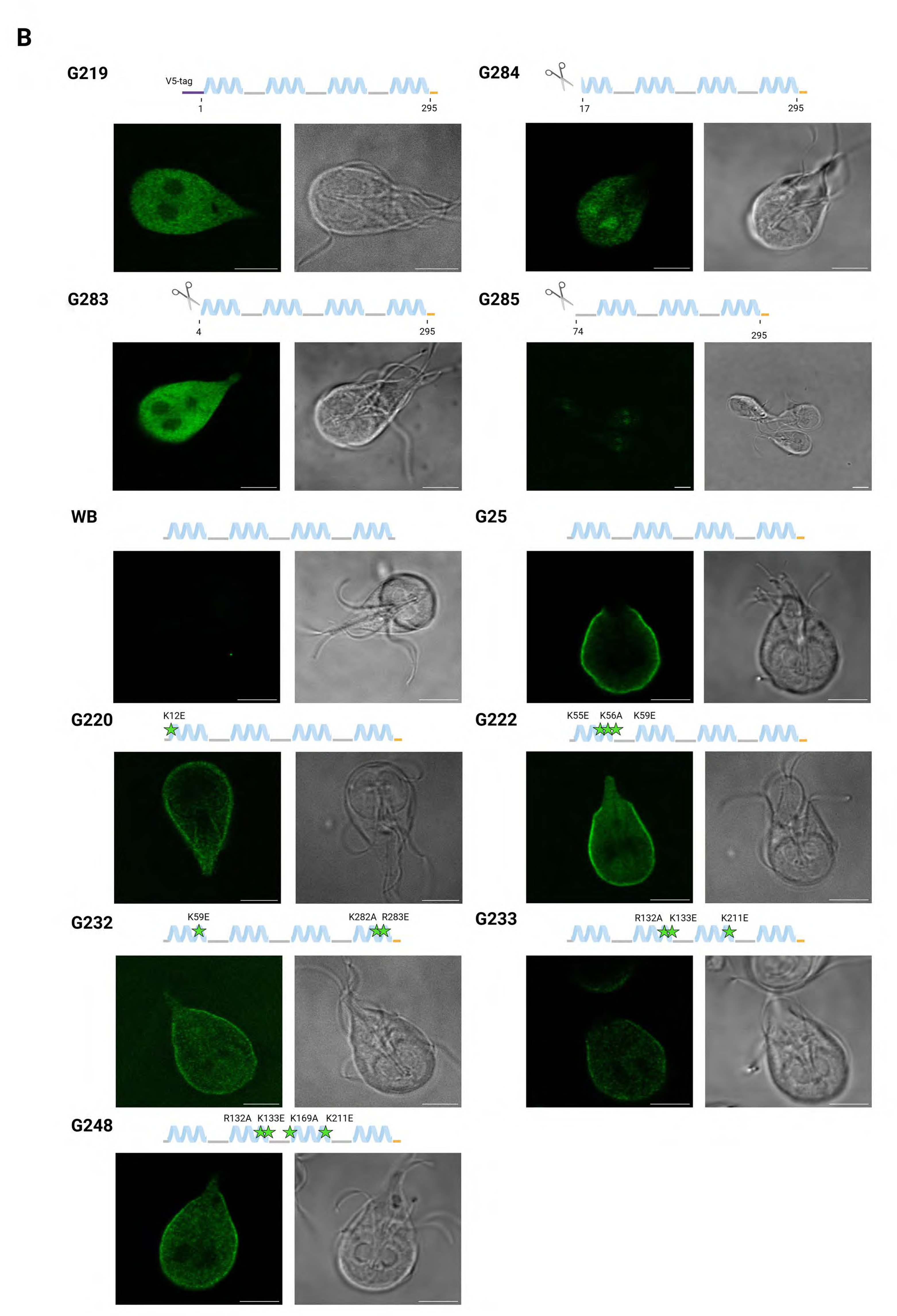
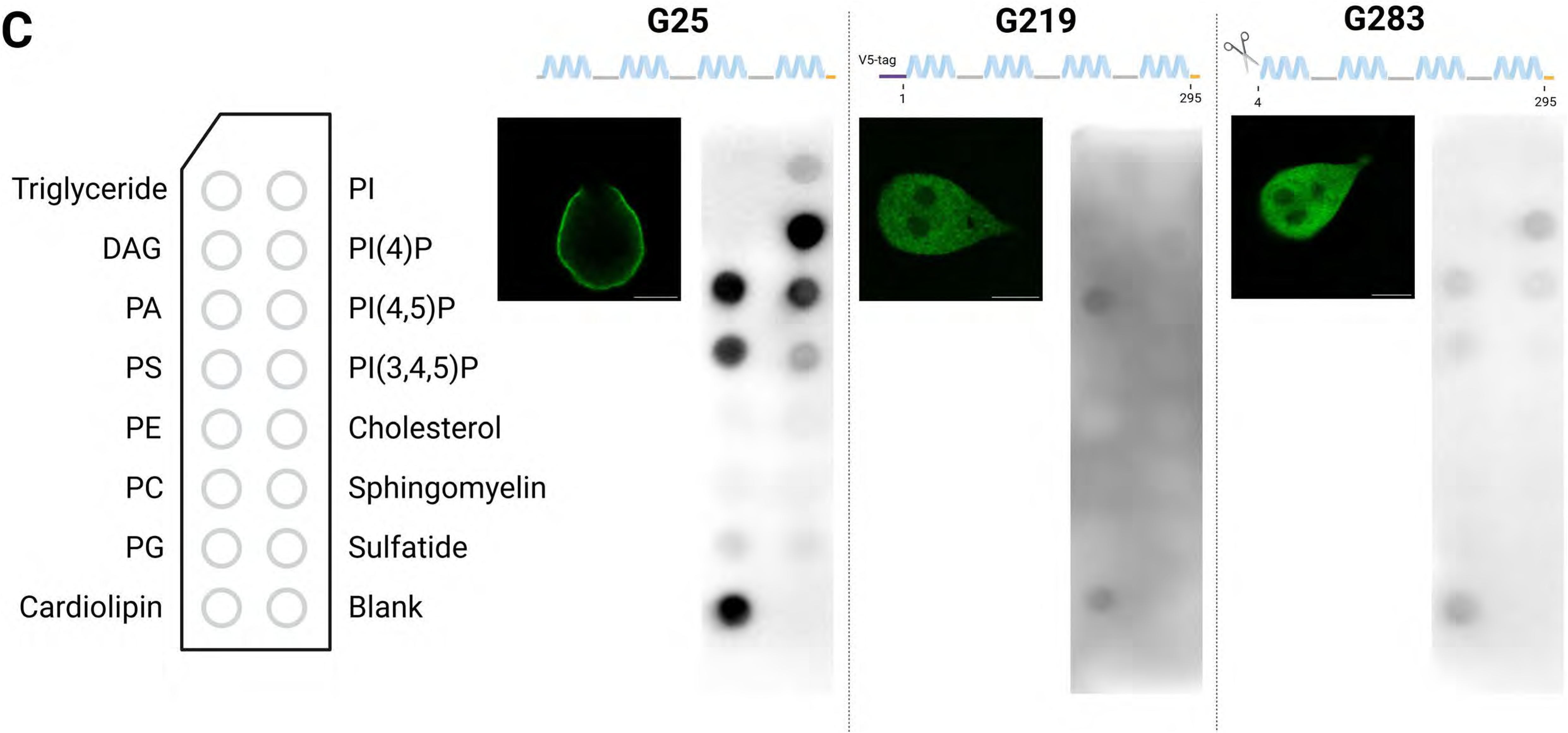
Site-directed mutagenesis and truncation analysis of α1G, compared to α8G and anxA5. **A.** Giardia lines G25 and G282 were developed to overexpress HA-tagged α1G, whereas line G455 was engineered to express codon-optimized human anxA5, similarly HA tagged at the C terminus. Lines G220, 222, 232, 233 and 248 were developed to express putative α1G lipid binding mutants, whereas lines G219, 283, 284 and 285 were engineered to express α1G N terminus variants, either by insertion of a V5 epitope tag, or by increasing truncations of the N terminal region. **B.** Confocal microscopy analysis following IFA of lines listed in A., including a non-transgenic WB control sample. **C.** Lipid strip binding data analysis for lines G25, G219 and G283, including confocal microscopy analysis of IFA samples for each line.

Giardia lines G25 and G282 (Figure 6A, tagged proteins) have hitherto been used to express wildtype HA-tagged α1G, and serve as reference-control lines. Lines G220, 222, 232, 233 and 248 (Figure 6A, positively-charged residues mutants) were engineered to express different combinations of Lys and Arg mutants which, based on XL-MS data modelling, were posited to play a role in lipid binding. On the other hand, lines G219, 283, 284 and 285 (Figure 6A, N-terminus mutants) were developed to interrogate the functional role of the short N-terminal region of α1G.

The outcome of the microscopy-based analysis of the expression of all listed constructs in the respective lines, shows that, of all the α1G variants tested, those with a modified N terminus, either by addition of a V5 epitope tag (line G219), or by deletion of progressively larger N-terminal segments, lost targeting to PECs (Figure 6B). PECs localization is already lost upon deletion of the first three amino acid residues (line G283). In line with these data, when HA-mediated co-IP eluates from lines G219 and G283 were tested on lipid strips, alongside the reference eluate from line G282, no appreciable lipid-binding activity could be detected (Figure 6C; Supplementary Data 3 and 4). All tested mutants for possible involvement in lipid binding were found at PECs, similar to reference constructs expressed in lines G25 and G282 (Figure 6B). To test each α1G variant in isolation as opposed to testing the behaviour of α1G-containing protein complexes extracted by co-IP from Giardia cell extracts, we attempted to express α1G in both *E. coli* and in Yeast. Although protein expression was overall successful (have to include in supp fig 1), both heterologous expression systems failed to provide functional protein (based on testing of lipid binding capacity on lipid strips; Supplementary Figure 1). Furthermore, to interrogate αG behaviour in living cells, we engineered constructs for expression of full-length α1G fused to mNeon-Green (mNG) at the C terminus (line G370), alongside expression of an mNG C-terminally tagged version of construct G283 (line G390) and development of line G391 for expression of an mNG C-terminally tagged α8G variant. A line engineered to express free mNG (line G369) was included as a control (Supplementary videos 1-4). Live cell imaging data show how, in contrast to mNG fusion of full-length α1G which associates to PECs and in line with previous data, the -3 deletion construct/(Δ3α1G (line G390) loses PEC targeting and mimics more closely the deposition of both a free mNG (line G369) and the mNG fusion to α8G (line G391).

### An intact N-terminus, no matter how short, is required for oligomerization of alpha1-giardin

Mass photometry (MP) was employed to assess the oligomeric state of protein samples under native conditions. The analysis included a control sample (WB) and four test proteins: G282 (α1G), G196 (α8G), and G283 (Δ3α1G) (Figure 7A). The WB sample showed a single, narrow peak (Figure 7A) corresponding to the background signal typically observed in mass photometry experiments (40-70 kDa).

**Figure 7.**
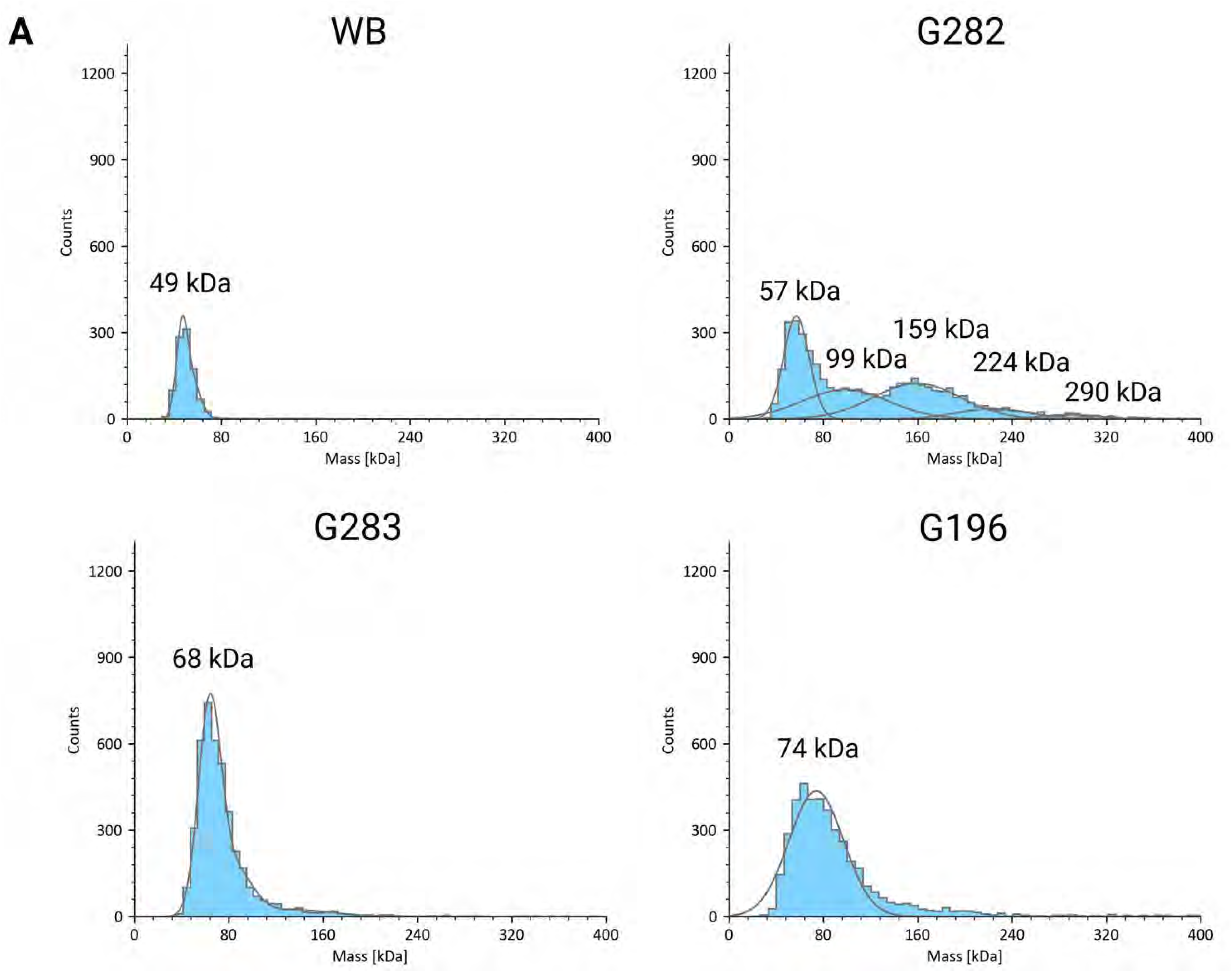
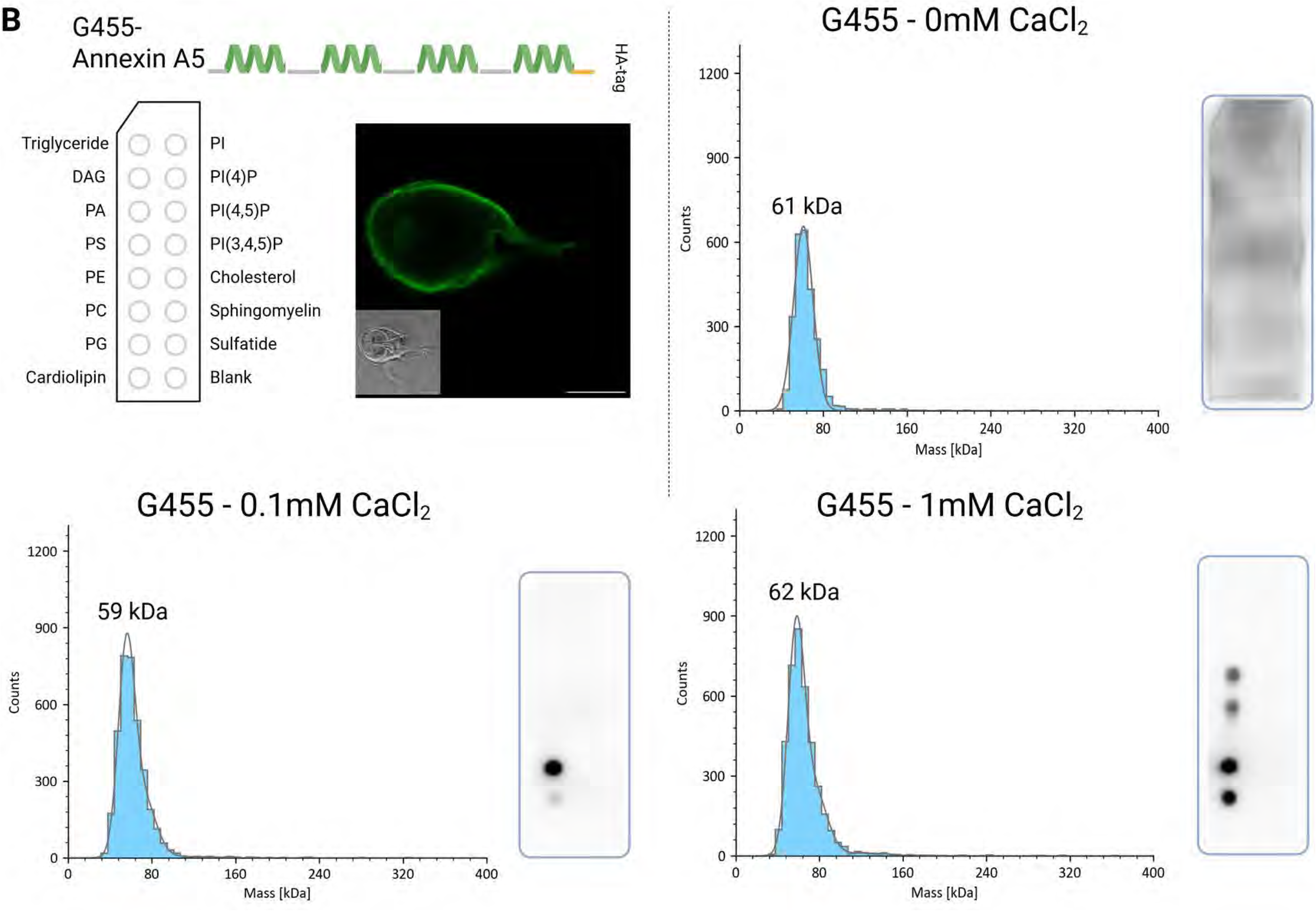
Mass photometry data analysis for HA-tagged α1G, Δ3α1G, α8G and anxA5, extracted from lines G282, G283, G196 and G455, respectively. **A.** Mass distribution (in kDa) of HA-tagged α1G, Δ3α1G, α8G containing complexes extracted from their respective lines (G282, G283 and G196) and resuspended in PBS, compared to a similarly prepped sample from non-transgenic WB cells as control. **B.** Mass distribution (in kDa) and lipid strip binding activity of Giardia-expressed anxA5 extracted from line G455 and resuspended in Hepes with increasing amounts of CaCl_2_. Please note that no lipids were added to this solution prior to both MP and lipid strip assays.

Interestingly, α1G, with a predicted monomeric mass of 35 kDa, showed a distinct pattern of multiple mass fractions, strongly indicating oligomer formation. In addition to the background peak, several higher-mass species were observed at approximately 99 kDa, 159 kDa, 224 kDa, and 290 kDa (Figure 7A). These values suggest the presence of oligomeric states, potentially corresponding to trimeric, pentameric, and higher-order assemblies. Although the higher-mass species may reflect complexes with α1G-binding partners, co-IP data indicate that α1G is the main component, with other proteins present only in trace amounts.

Along with α1G, we tested α8G as a representative cytosolic giardin and the truncated form of α1G - G283. Both proteins extracted from G196 and G283 displayed only a single peak centered around 70 kDa (Figure 7A). No higher-order mass species were observed in either sample. Although the formation of dimers could theoretically occur, such species would fall below the detection threshold of the instrument. Therefore, these samples are interpreted as being in a monomeric form, under the conditions tested, with no detectable oligomerization.

### Giardia-expressed human annexin A5 localizes to PECs and binds positively-charged lipids

Unlike parasitic annexins, which are under-investigated, mammalian annexins have been reported on extensively. One of the better functionally and structurally characterized annexins, which was also developed as a molecular detection tool for pre-apoptotic cells, is human annexin A5 (anxA5) (Huber *et al*., 1990; Chen, Sheldon and Pincus, 1993; Demange *et al*., 1994; Trotter, Orchard and Walker, 1995; Sopkova *et al*., 1998; Campos *et al*., 1999; Oling, Bergsma-Schutter and Brisson, 2001; Mo *et al*., 2003; Patel *et al*., 2005; Bouter *et al*., 2011a; Zhu *et al*., 2014; Lin, Chipot and Scheuring, 2020a; Lichocka *et al*., 2022a). AnxA5 is known to require calcium ions and oligomerization for lipid binding. We wondered whether anxA5 expressed in Giardia would behave similar to α1G in terms of PEC targeting and lipid binding properties. To address this question, we engineered line G455 to express HA-tagged Giardia codon-optimized anxA5. Confocal microscopy analysis following IFA showed deposition of the HA-tagged reporter at the periphery of the cell, clearly proximal to PECs (Figure 7B). Giardia-expressed anxA5 was further extracted for testing in lipid strip assays at different calcium concentrations. In the absence of calcium, there was no detectable lipid binding (Figure 7B). As calcium levels increased, we detected binding to Cardiolipin and Phosphatidylglycerol at 0.1 mM Ca²⁺, and additional binding to Phosphatidic acid and Phosphatidylserine at 1 mM Ca²⁺. The same anxA5 protein sample was analysed by MP. No oligomer formation could be detected in the MP assay under any condition. However, the reported trimerization of human anxA5 (Bouter *et al*., 2011b) depends on lipid binding (Richter *et al*., 2005), and no lipids were included in the MP assay.

### Molecular dynamics of oligomeric alpha1-giardin variants and membrane binding simulations

To investigate the membrane-binding behaviour of the α1G trimer, molecular dynamics simulations were performed across five distinct lipid compositions (Supplementary Table 6). The base membrane contained phosphatidylcholine (PC), phosphatidylserine (PS), and cholesterol, while the other systems included either 3% or 12% of PI3P or PI5P (Supplementary videos 5-24). In all cases, the protein interacted with the membrane, but the nature of the interaction changed depending on the presence, type and amount of PIP. In the case of the basic membrane, the interaction was short-lived, and the trimer dissociates quickly. However, after the addition of PIPs, the interaction was stabilized.

Average values for membrane interactions were calculated using residues with k*_off_* bootstrap averages not exceeding 0.5 ns⁻¹. Membranes containing PIPs demonstrated enhanced interaction characteristics compared to the basic membrane. On average, the trimer bound to PIP-containing membranes stayed longer, with six times the duration, seven times the residence time, and five times the occupancy compared to PS-only membranes. Based on a lower k*_off_*, it also detached four times more slowly, as shown in Figure 8A. This shows increased affinity to membranes containing PIPs in comparison to the PS-only membrane, highlighting the importance of PIPs in stabilizing α1G at the membrane interface. These findings suggest a stronger affinity of α1G for PIP-enriched membranes, which is in line with experimental data and highlights the role of PIPs in stabilizing α1G at the membrane interface.

**Figure 8.**
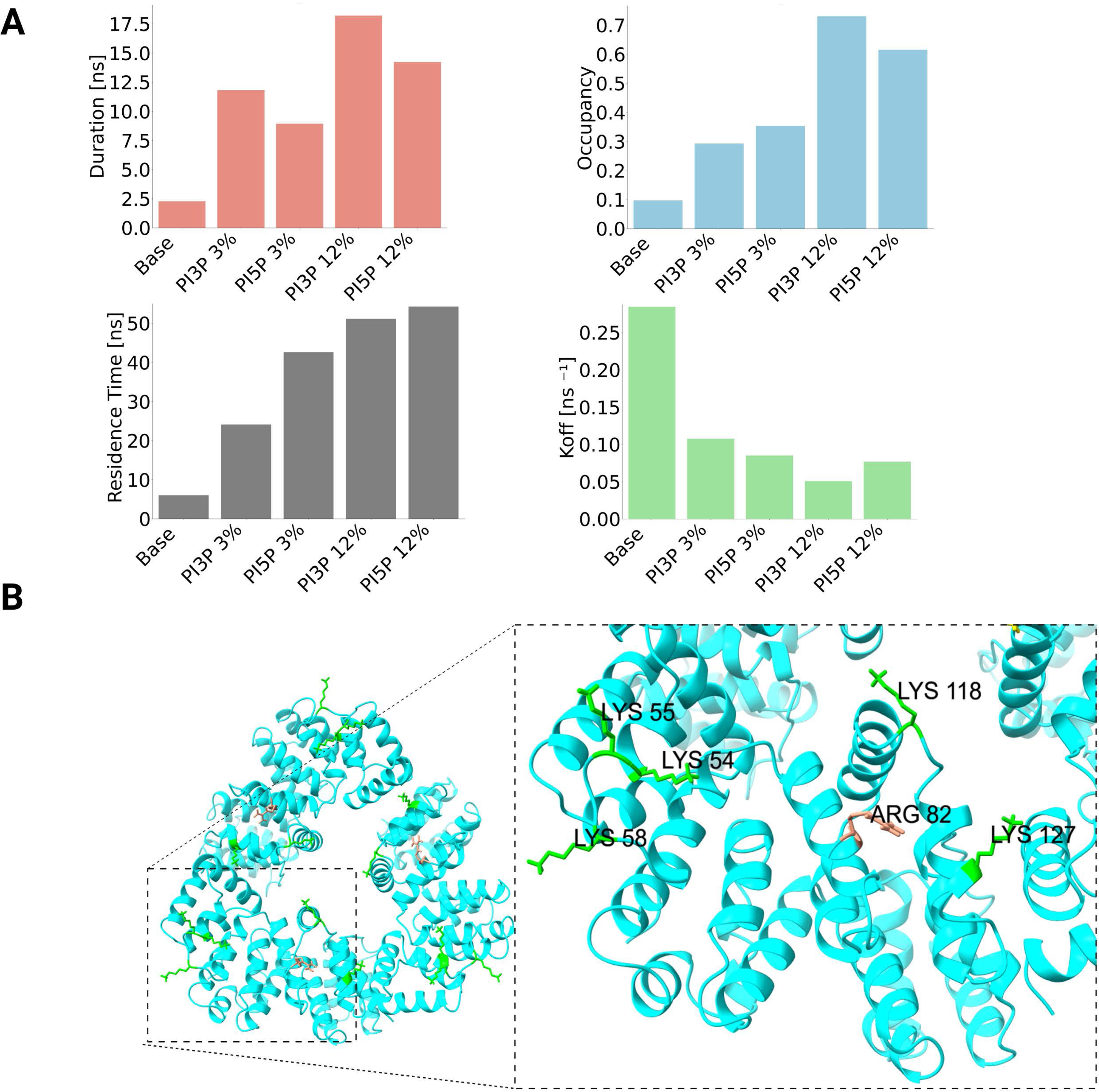
Molecular dynamics simulations. **A.** Average values, calculated with PyLipID, for duration, occupancy, residence time and k*_off_* for the a1g trimer interacting with different enrichment levels for PI3P and PI5P at target membranes during MD simulations. **B.** Lysine (lys) and arginine (arg) residues on the a1g trimer (cyan) involved in membrane contacts: dark tuna highlights show stable, long-lasting interactions, while green marks frequent but short-lived contacts.

Among the PIP variants, the stability of interaction increased with increasing amounts of PIPs for both PI3P and PI5P. 12 % PI3P membrane showed, on average, the longest duration of single interactions and the highest occupancy with the smallest k*_off_* value (Figure 8A). The longest average residence time was observed for 12% PI5P. Surprisingly, for a smaller concentration of PIPs, PI3P outperformed PI5P only in duration time, whereas PI5P consistently outperformed PI3P across other parameters, including occupancy, residence time, and dissociation rate.

To better understand which regions of α1G contribute to lipid binding, protein-lipid interaction parameters across all simulations were analysed (Supplementary Data 6 and Supplementary Figure 2). Based on duration, occupancy, k*_off_*, and residence time, the most significant interacting residues were identified and grouped into two categories (Figure 8B). The first group comprises residues that form stable, long-lived contacts—defined as interactions lasting over 50 ns, with residence times above 50 ns and very low dissociation rates (k*_off_* < 0.01 ns⁻¹). Only Arg82 met all these criteria, and only in the 12% PI3P membrane variant. In contrast, Arg82 did not meet the threshold for the PI5P variant, falling short due to a slightly lower duration (∼35 ns). Notably, the occupancy of Arg82 is moderate, which is in line with rarer but more stable interactions.

The second group consists of residues that interact more transiently but frequently. These residues exhibited occupancy values above 0.5, shorter contact durations (under 20 ns), residence times between 50 and 200 ns, and k*_off_* values below 0.05 ns⁻¹. Residues meeting these criteria were identified only in the 12% PIP lipid variants. For PI3P, four residues were selected—Lys127, Lys55, Lys118, and Lys58—while for PI5P, only two residues met the criteria: Lys55 and Lys54 (Figure 8B). Notably, more residues within the trimer interact with PI3P, yet these interactions tend to have shorter durations compared to PI5P. Additionally, PI5P interactions show lower k*_off_* values and higher residence times, suggesting that although fewer residues are involved, their interactions may be more stable and functionally significant.

## Discussion

The data presented in this report point to a fundamental role for the N-terminus of α1G in oligomerization, lipid-binding ability and correct subcellular targeting to PECs. To synthesize all the data presented so far, while taking stock of previous reports (Weiland *et al*., 2003b; Wei *et al*., 2010b; Weeratunga *et al*., 2012b; Einarsson *et al*., 2016b; Ma’ayeh *et al*., 2017; Davids *et al*., 2019; Balmer, Wirdnam and Faso, 2023; Warmus, Balmer and Faso, 2025), we propose a new working model for αGs at PECs, with a focus on α1G as an immunodominant and highly promising Giardiasis vaccine candidate (Weiland *et al*., 2003c), implicated in UPS mechanisms at the parasite-host interface (Balmer, Wirdnam and Faso, 2023) (Figure 9).

**Figure 9.**
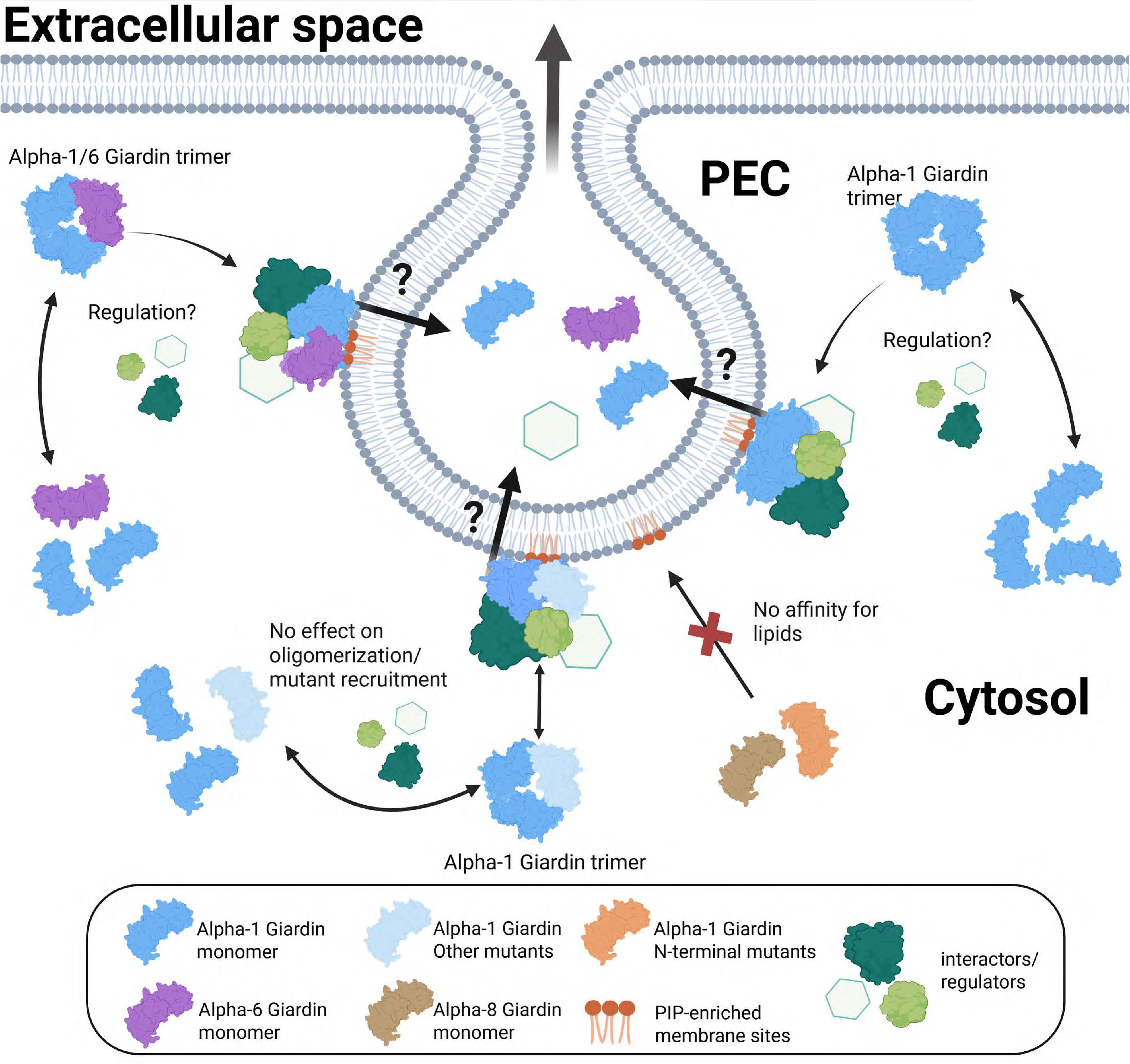
A new working model for PECs-associated and diffused alpha-Giardins. α1G (dark blue) and α6G (violet) are shown as examples of PECs-associated, whereas α8G (brown) is used as an example of a cytosolic alpha-giardin. Compared to α8G which does not present detectable lipid-binding activity, both α1G and α6G bind lipids and localize to PECs. The XL-MS data point to the existence of both homo and hetero-dimers of α1G and α6G. α1G putative lipid binding mutants (light blue) are still detected at PECs. However, α1G N-terminal variants (burnt orange; either deletion or V5-tagged mutants) mimic α8G in the absence of both PECs localization and lipid-binding activity. This indicates how the N-terminal region of α1G is an essential functional domain, likely linked to the need for oligomerization to enable lipid binding capacity. These observations are corroborated by MP data presented in this report, showing how α1G N-terminal variants remain monomeric in solution, unlike the wt α1G control. Several open questions remain, including the role for interaction partners previously identified at PECs-associated complexes (green components) as regulators of oligomerization, followed by lipid binding and mediation of UPS at PECs (Balmer, Wirdnam and Faso, 2023).

We postulate that αGs at PECs such as α1G (Figure 9, dark blue) and α6G (Figure 9, violet), unlike αGs which are never recruited to these organelles, can oligomerize and, consequently, can bind specific lipid residues at PECs. Oligomerization, and therefore lipid-binding ability, is dependent on an intact N-terminus which is therefore also ultimately responsible for PEC localization. In contrast, α8G (Figure 9, brown) is never found at PECs, nor does it detectably bind lipids, nor does it manifest any evidence for oligomerization. Unlike putative α1G lipid-binding mutants (Figure 9, light blue) which are indistinguishable from wildtype α1G, N-terminal α1G mutants (Figure 9, orange) behave similarly to α8G, further corroborating the essentiality of α1G’s N terminal region for oligomerization, lipid binding activity, and deposition at PECs.

Several open questions remain, most notably the identity of the regulators of αG oligomerization (Figure 9, green components) as a pre-requisite for lipid binding at PECs, including a role for αGs at PECs in mediating UPS of selected cargo (Figure 9, green components).

### Why does heterologous expression of α1G fail to give lipid-binding protein?

An immediate question arising from the working model proposed above concerns the nature of the regulatory circuits allowing some αGs to associate to PEC membranes, while others, although virtually indistinguishable on a sequence level, appear not to. Since the entire family of αGs has not yet been systematically tested for this property, we cannot claim oligomerization as exclusive to PECs-associated αGs. However, MP and XL-MS data do point towards a major role for oligomerization in functionality and PECs association, at least for α1G. At the same time, failure to express lipid binding-competent α1G in heterologous systems, both prokaryotic and eukaryotic (Supplementary Figure 1), points to a missing essential Giardia-specific component to obtain functional protein. This component may either be a missing yet essential interaction partner or a dedicated post-translational modification (Figure 9, green components). The MS data point to the presence of several phosphorylated residues which may be integral to protein function and may not be recapitulated in heterologous systems in the absence of a specific interaction partner. One such candidate partner regulator could be NEK kinase-GL50803_15411 which, although still classified as a catalytically inactive kinase (as the majority of Giardia NEKs) (Manning *et al*., 2011), may act as a scaffold for other (active) kinases in Giardia. In a previous study, N-terminally tagged α1G expressed in *E. coli* was reported to successfully bind glycosaminoglycan (Weiland *et al*., 2003b). This report is compatible with our data while also showing how the two ends of this protein perform different and independent functions which are differentially impacted by attempts at heterologous expression.

### A comparison of α1G and *Giardia*-expressed anxA5

Epitope-tagged anxA5 expressed in Giardia localizes to the periphery of the cell, consistent with PEC association and lipid-binding activity of extracted HA-anxA5. This corroborates the observation that only αGs at PECs can bind lipids, as also shown using lipid strip experiments. AnxA5 was shown to organize in trimers at the membrane (Lin, Chipot and Scheuring, 2020b), and anxA5 oligomerization depends on PTMs, negatively charged lipids and calcium concentration (Lichocka *et al*., 2022b). This last variable is perhaps the most conspicuous difference between α1G and anxA5, where the former’s lipid binding ability was never impacted by the absence of calcium ions, whereas the latter clearly depends on them. In this sense, α1G is an unusual lipid-binding annexin at unusual endocytic compartments, with unusual requirements for calcium (Weeratunga *et al*., 2012b). However, the common need for oligomerization is a shared feature which points once more to the importance of this structural property as a pre-requisite for membrane binding.

## Conclusion

In conclusion, the data presented highlights the critical role of the N-terminus of α1G in its oligomerization, lipid-binding ability, and correct localization to PECs, positioning it as a promising Giardiasis vaccine candidate. The proposed model suggests that αGs like α1G and α6G, but not α8G, require an intact N-terminus for effective oligomerization and lipid interaction at PECs. Moreover, the inability to express lipid-binding competent α1G in heterologous systems points to a potential Giardia-specific component that is crucial for its functionality, possibly involving specific post-translational modifications or interaction partners. The comparison with anxA5 further emphasizes the importance of oligomerization in membrane binding, highlighting unique characteristics of α1G in lipid binding at PECs. The data underscore the complex and essential role of α1G in the parasite’s biology; any insights in its structural and functional determinants, as presented in this report, provide additional information towards the fulfilment of α1G’s potential as a highly promising Giardiasis vaccine candidate

## Materials and Methods

### Construct generation, transformation and cell culture conditions

The DNA constructs used in this study were generated by PCR, followed by ligation into expression vectors using either T4 DNA ligase or Gibson assembly. The sequences of the oligonucleotides and all the details of the reactions are summarized in Supplementary Data 5.

*Giardia* trophozoites were cultured in previously well-described axenic conditions (Cernikova, Faso and Hehl, 2020; Santos *et al*., 2022; Balmer, Wirdnam and Faso, 2023). Cells were transfected via electroporation with circular plasmid vectors and selected with the antibiotic puromycin. Immuno-blotting as well as IFAs (immunofluorescence assays) were used to check expression levels of the transfected constructs.

### Immunofluorescence assay

IFAs were performed according to previously published protocols (Santos *et al*., 2022; Balmer, Wirdnam and Faso, 2023). Trophozoites, wild type or expressing the epitope tagged protein of interest, were grown in 12ml Nunc polystyrene culture tubes (ThermoFisher Scientific). Per IFA one such tube per line was grown until the cells were forming a confluent layer. The tubes were then incubated on ice for a minimum of 30 min to detach the cells. The cells were then pelleted at 900g for 10 min, washed in PBS (phosphate-buffered saline) and fixed in 3% formaldehyde (Sigma) in PBS for a minimum of one hour or overnight. Fixation was followed by quenching in 0.1M Glycine in PBS and permeabilization in 2% bovine serum albumin (BSA) + 0.2% Triton-X-100 in PBS. For antibody staining the cells were first incubated for one hour in 2% BSA + 0.1% Triton-X-100 in PBS containing the rat-derived monoclonal primary anti-HA antibody (Roche) diluted 1:250. Then the samples were washed twice in 1% BSA + 0.05% Triton-X-100 in PBS (wash solution). The secondary antibody was diluted 1:250 in 2% BSA + 0.1% Triton-X-100 in PBS. A goat-derived anti-rat IgG (H+L) conjugated to Alexa Fluor 488 (ThermoFisher) was used for incubation in the dark at room temperature. After the second incubation cells were again washed twice in wash solution and then resuspended in Vectashield embedding medium (Reactolab) containing DAPI (4′-6-diamidino-2-phenylindole) as a DNA label. Imaging was performed at a Leica SP8 confocal microscope configured with white light lasers.

### Co-immunoprecipitation (co-IP) in native conditions and DSSO crosslinking

Native co-IPs were performed as previously described (Balmer, Wirdnam and Faso, 2023). *Giardia* trophozoites expressing a tagged reporter construct of interest or wild type control WB were grown in two T25 culture flask per line over two days until confluency was reached on both sides of the flask (turned after the first 24 hours). For harvesting, the cultures were placed on ice for a minimum of one hour for the cells to detach before they were transferred to 50ml Falcon tubes where the cells were resuspended in 10ml PBS with addition of protease inhibitors (Sigma) and 1% Triton-X-100. Cells were burst using sonication and then incubated in the cold for two hours with gentle rotation. Samples were transferred to Eppendorf tubes and centrifuged at 16000g for 10 min at 4°. Supernatants were filtered through 0.2μm PVDF membrane syringe filters and incubated with HA agarose beads (Thermo Fisher Scientifc) for protein pulldown according to manufacturer’s instructions (40µl slurry/extract from a T25 flask). Following the overnight incubation, the beads were washed 3x with PBS+ 0.1% Triton-X-100 and 3x with PBS. For simple co-IP, beads were stored frozen before analysis.

For DSSO crosslinking, beads corresponding to one flask per line were resuspended in 500ul PBS and 10ul DSSO in DMSO (50mM) was added to reach approximately a 100-fold molar excess of crosslinker over the protein concentration. After an incubation of 45 minutes at room temperature, the reaction was stopped with Tris buffer (resulting in a final Tris concentration of 0.1M) for 5 min and the beads were washed 3x with PBS and stored frozen before analysis. All bead samples were handed over to the Core Facility for Proteomics and Mass Spectrometry of the University of Bern for tryptic digests and peptide analysis.

### Liquid Chromatography Mass Spectrometry for native and DSSO crosslinked samples for XL-MS

Proteins captured on the pull-down beads were denatured in 30 μL 8 M urea, 50 mM Tris/HCl pH 8, and cysteines alkylated with 50 mM iodoacetamide for 30 min in the dark after reduction with 10 mM DTT for 30 min at 37°C. The urea concentration was lowered by adding 60 μL 20mM Tris/HCl pH 8 containing 2 mM CaCl_2_, followed by digestion with 100 ng LysC for 2 hours at 37°C. Urea was further diluted to 1.6 M and digestion continued by addition of 100 ng trypsin over night at room temperature. After acidification with 1% TFA end concentration, aliquots of 5 μL were analyzed by nano-liquid reversed phase chromatography coupled to tandem mass spectrometry on an Orbitrap Fusion LUMOS mass spectrometer that was coupled with a Dionex Ultimate 3000 nano-UPLC system (ThermoFischer Scientific, Reinach, Switzerland). For non-crosslinked samples a standard data-dependent acquisition method as described elsewhere (Buchs et al. 2018) was used with a homemade C18 CSH Waters separation column (1.7 μm, 130 Å, 75 μm × 20 cm) at a flow rate of 250 nL/min. The cross-linked samples were analyzed on the same system using a MS2-MS3-MS2 method. Full MS scans in the orbitrap in the range of 375-1500 m/z were acquired every 5 s. In between data-dependent CID MS2 scans on peptide ions with charge 3-8 were acquired in the orbitrap with resolution 30’000, a fixed relative collision energy of 25%, precursor isolation window of 1.6 m/z, and a MIT of 54 ms. Two MS3 CID fragmentations were triggered on MS2 fragments which formed an ion pair spaced by the DSSO-specific 31.9721 Da at a mass tolerance of 10ppm. Relative collision energy was set at 35%, normalized automatic gain control to 200%, and MIT to 120 ms. If no ion pair was detected in the MS2 CID spectrum another MS2 scan with HCD fragmentation was recorded in the orbitrap at a relative collision energy of 30% and MIT of 100 ms. The non-cross-linked data was interpreted with MaxQuant (version 2.3.1.0) applying strict trypsin cleavage rule with maximum 3 missed cleavages, carbamidomethylation of cysteines as fixed, oxidation of methionines and protein n-terminal acetylation as variable modifications allowing a maximum of three modifications per peptide, a first round search precursor mass tolerance of 15 ppm, and fragment mass tolerance of 0.4 Da, against the *Giardia* intestinalis Assemblage AWB protein sequence database from *Giardia*DB (version 55) together with a small database of 230 proteins commonly found as contaminants and the reversed versions of both databases. A false discovery rate on the peptide and protein level of 1% was applied and only proteins identified with at least 2 unique peptides were accepted. The data of the crosslinked samples was interpreted against the same protein sequence database using ProteomeDiscoverer version 3.0 SP1 with built-in Sequest HT and XlinkX again using a decoy search strategy to control the false discovery rate at 1% for non- and cross-linked peptide identifications.

### Lipid strip protein binding assay

Proteins were obtained by anti-HA-agarose beads pulldown as described in the co-IP section. Afterwards, the beads were eluted with 2mg/ml HA peptide (MedChemExpress) in PBS.

Membranes spotted with a selection of lipids (Echelon Biosciences P-6002) were used for lipid binding assays according to manufacturer’s protocols with each of several alpha giardins prepared as described above. The membranes were incubated in water for 5 minutes followed by a 1h blocking step in PBS containing 3% lipid-free BSA (Sigma). Then alpha giardins were added and incubated for 1 h. Membranes were washed 3×10min with PBS/Tween 0.05% and then incubated for 1h in PBS containing primary antibody (rat-derived monoclonal anti-HA, Roche). Membranes were washed again, incubated in secondary antibody (goat anti rat coupled to HRP, Sigma) before a last set of washes. Membranes were then imaged using enhanced chemiluminescence (ECL, ThermoScientific).

### Structural reconstruction using XL-MS data

XL-MS data was filtered for identified intra- and interlinks with a threshold of Delta XlinkX Score ≥ 50, using the Delta XlinkX Score as a proxy for hit-reliability. An interactome network was generated using the Cytoscape software based on found interlinks. Found interlinks between alpha-1 giardin and alpha-6 giardin and Xaa-pro dipeptidase in the XL-MS data were visualised as linear maps informed by data on protein motifs from *Giardia*DB. To reconstruct the 3D structure of the identified associated proteins alpha-1- and alpha-6 giardin and Xaa-pro dipeptidase, structural models were sourced from alphafold.ebi.ac.uk or predicted using AlphaFold webserver(Abramson *et al*., 2024). The HADDOCK 2.4 web server interface(Dominguez, Boelens and Bonvin, 2003; Van Zundert *et al*., 2016) was then used to integrate our own generated post-capture XL-MS data with the sourced 3D models of the individual protein structures to construct a comprehensive model of the 3D structure of these three interactors. The resulting structure was subsequently refined and merged using ChimeraX (Pettersen *et al*., 2021), GIMP (The GIMP Development Team 2019, no date) and Cytoscape (Shannon *et al*., 2003).

### Mass photometry

Mass photometry experiments were conducted using a Refeyn TwoMP system (Refeyn Ltd., Oxford, UK) equipped with the AcquireMP and DiscoverMP software packages for data acquisition and analysis, respectively, under standard settings. For the experiments, high-precision microscope coverslips, sourced from Refeyn, were used as single-use items. To preserve the droplet shape of the sample, self-adhesive silicone culture wells (Grace Bio-Labs reusable CultureWell™ gaskets) were employed. Alpha giardin samples and WB were diluted in PBS, while annexin A5 samples were prepared in 20 mM HEPES (pH 7.5), 300 mM NaCl, and the appropriate concentration of CaCl₂.

For contrast-to-mass calibration, Bovine Serum Albumin Fraction V Low Heavy Metals (Millipore) oligomers with molecular weights of 66, 132 and 198 kDa were utilized. Prior to measurements, protein stocks were diluted in stock buffers containing 20 mM HEPES (pH 7.5) and 300 mM NaCl for Annexin A5 experiments or PBS for alpha giardins measurements. Specifically, 2 µL of the protein solution was mixed with 18 µL of analysis buffer, yielding a final droplet volume of 20 µL with a protein concentration of approximately 1 µg/mL.

### Protein Modelling and Initial Structure Preparation

To investigate the interaction between α-1-giardin and lipid bilayers, a trimeric model of α-1-giardin was constructed. The monomeric structure was predicted using the AlphaFold web server(Abramson *et al*., 2024), excluding the N-terminal methionine (Met1) to reflect the mature protein form. Based on XL-MS data identifying interaction sites, a trimeric configuration was assembled using the HADDOCK v2.4 docking platform(Dominguez, Boelens and Bonvin, 2003; Van Zundert *et al*., 2016). The trimer was selected for simulation based on evidence from mass spectrometry indicating its oligomeric state.

Five distinct membrane systems were constructed using the CHARMM-GUI Membrane Builder webserver(Mehnert *et al*., 2006; Jo, Kim and Im, 2007; Jo *et al*., 2008, 2009; Brooks *et al*., 2009; Wu *et al*., 2014; Lee *et al*., 2019; Park *et al*., 2021; Feng *et al*., 2023; Brown *et al*., 2024; Gee *et al*., 2024), with each consisting of approximately 350 lipid molecules per leaflet. All systems included PI(3)P and PI(5)P lipids at varying concentrations to assess their influence on alpha-1 giardin binding. The exact lipid compositions for each system are summarized in Supplementary Data 6. Water molecules were removed from the unit cell, and the α-1-giardin trimer was positioned above the membrane at a minimal distance of 10 Å from the surface to allow for potential interaction while avoiding initial steric clashes.

### Molecular dynamics simulations

The membrane system preparation was continued in CHARMM-GUI, with all components parameterized using the CHARMM36 force field, generating GROMACS-compatible files(Mehnert *et al*., 2006; Jo, Kim and Im, 2007; Jo *et al*., 2008, 2009; Brooks *et al*., 2009; Wu *et al*., 2014; Lee *et al*., 2019; Park *et al*., 2021; Feng *et al*., 2023; Brown *et al*., 2024; Gee *et al*., 2024).

The TIP3P water model was used to solvate each system in a simulation box of averaged dimensions 170 × 140 × 140 Å, containing approximately 350000 atoms. Appropriate numbers of Na⁺ and Cl⁻ counterions were added to ensure electrostatic neutrality and physiological ionic strength (∼0.15 mM).

Energy minimization was performed using the steepest descent algorithm, with a convergence Vmax of 1000 kJ·mol⁻¹·nm⁻¹. Partial equilibration was conducted on the CHARMM GUI server using the NVT and NPT ensembles to gradually relax positional restraints on the thermos-coupled components of the systems. Electrostatic interactions were treated using the particle mesh Ewald (PME) method, while van der Waals forces were computed via a force-switch cutoff scheme with a switching distance of 1.0 nm and a cutoff of 1.2 nm. Temperature coupling was applied separately to the solute, membrane, and solvent groups using the V-rescale thermostat with a reference temperature of 310 K and a coupling constant of 1.0 ps. Pressure was maintained at 1 bar using the C-rescale barostat with semi-isotropic coupling, a compressibility of 4.5×10⁻⁵ bar⁻¹, and a relaxation time of 5.0 ps. Finally, maintaining the same barostat, production runs of 200 ns were carried out using GROMACS 2023.3 (Abraham *et al*., 2023) with a leap-frog integrator and a time step of 2 fs, applying the LINCS algorithm to constrain all hydrogen-containing bonds. All simulations, including the final equilibration and production runs, were performed using 1 CPU core and GPU RTX4090 of University of Bern cluster UBELIX(no date). Protein-lipid analysis was performed with the Python library PyLipID (Song *et al*., 2021) with a dual cutoff of 0.15 nm – 0.25nm.

## Supplementary Files

**Supplementary Figure 1. Heterologous expression tests for epitope-tagged α1G in *E. coli* and Yeast**. Both systems yield sufficient protein however, lipid binding functionality is not maintained.

**Supplementary Figure 2. All raw data for MD simulation experiments.**

**Supplementary Data 1**_alpha1_6_11_8_native

**Supplementary Data 2**_XL_MS_intra-inter a-1,6,8,11-g

**Supplementary Data 3**_ Native IP for G282 and G283

**Supplementary Data 4**_XL-MS data for lines G282 and G283

**Supplementary Data 5**_Oligonucleotides

**Supplementary Data 6**_Protein-Lipid interaction simulation data

**Supplementary video 1**

**Supplementary video 2**

**Supplementary video 3**

**Supplementary video 4**

**Supplementary video 5**

**Supplementary video 6**

**Supplementary video 7**

**Supplementary video 8**

**Supplementary video 9**

**Supplementary video 10**

**Supplementary video 11**

**Supplementary video 12**

**Supplementary video 13**

**Supplementary video 14**

**Supplementary video 15**

**Supplementary video 16**

**Supplementary video 17**

**Supplementary video 18**

**Supplementary video 19**

**Supplementary video 20**

**Supplementary video 21**

**Supplementary video 22**

**Supplementary video 23**

**Supplementary video 24**

**Supplementary video 25**

## Acknowledgements

This work was supported by Swiss National Science Foundation grants PR00P3_179813 and 310030_219372, awarded to C.F. D.W. benefited from a short research stay funded by GHA-SciEx, the scientific exchanges funding scheme of the Africa-Europe Cluster of Research Excellence-Genomics for Health in Africa (CoRE-GHA). O.T-B and C.F. are members of CoRE-GHA, Genomics for Health in Africa (GHA), an Africa-Europe Cluster of Research Excellence. We are grateful to the PMSCF and the MIC for their service as core facilities at the University of Bern. Furthermore, we are grateful to Dr. Yoan Duhoo, Dr. Florence Pojer, and the team at the Protein Production and Structure Core Facility-EPFL (Switzerland) for their expert guidance in protein purification via FPLC, as well as their assistance with the acquisition and interpretation of mass photometry data.

## Author contributions

**Dawid Warmus:** Conceptualization, Data Curation, Formal Analysis, Methodology, Investigation, Validation, Visualization, Funding Acquisition, Writing – Original Draft Preparation, Writing – Review & Editing Preparation

**Corina D. Wirdnam:** Conceptualization, Data Curation, Formal Analysis, Methodology, Investigation, Validation, Writing – Original Draft Preparation

**Erina A. Balmer:** Data Curation, Formal Analysis, Methodology, Investigation

**Tendai Muronzi:** Supervision, Data Curation, Formal Analysis, Methodology

**Manfred Heller:** Data Curation, Formal Analysis, Methodology

**Özlem Tastan Bishop:** Resources, Supervision

**Carmen Faso:** Conceptualization, Funding Acquisition, Methodology, Project Administration, Supervision, Visualization, Writing – Original Draft Preparation, Writing – Review & Editing Preparation

