## Supplementary figures and images for "The N’ terminus of Alpha-1 Giardin, a parasitic annexin orthologue, is essential for oligomerization and lipid-binding activity"

### Supplementary Figure 1_new.jpeg

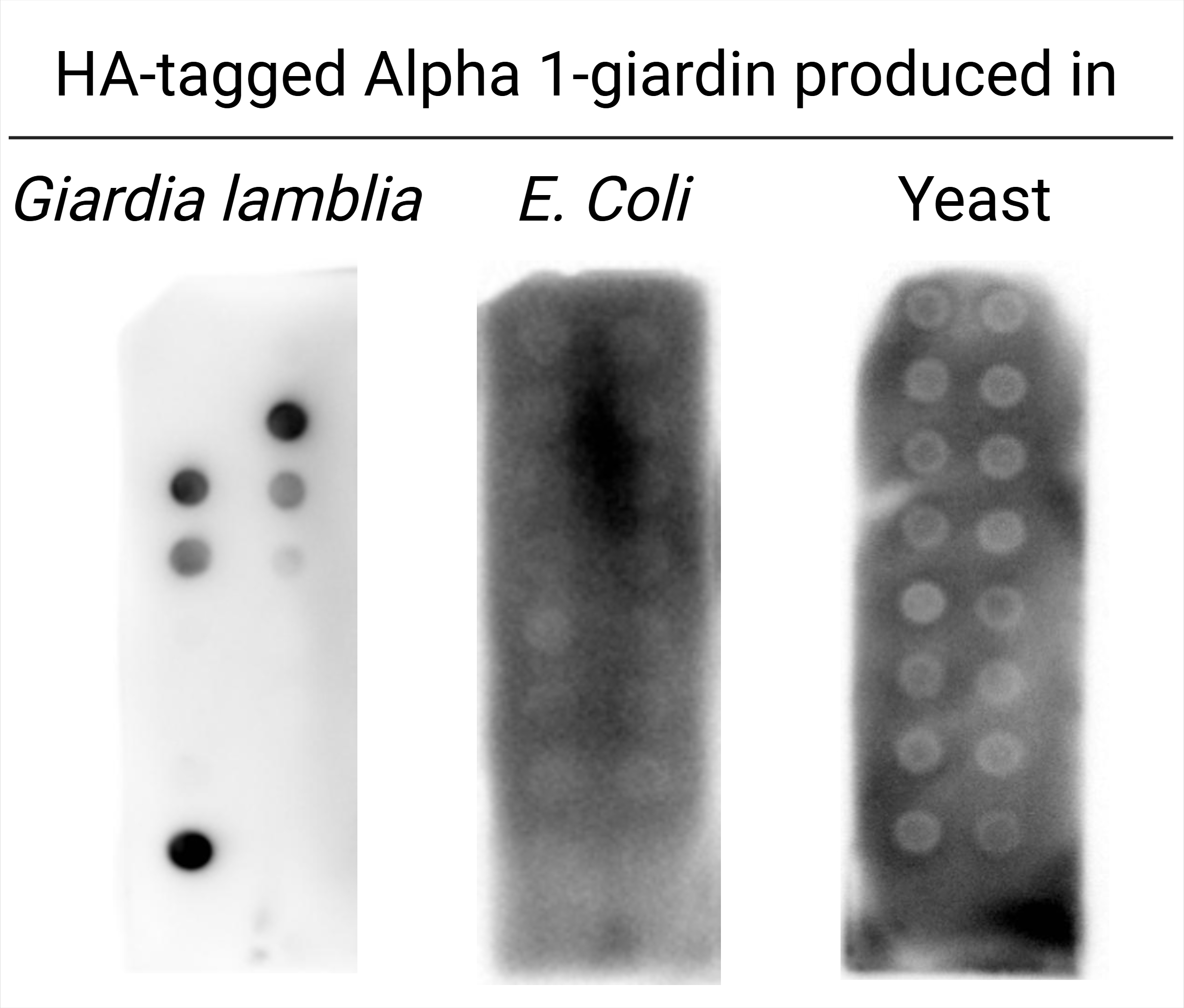
